# A stochastic hybrid model of DNA replication incorporating 3D protein mobility dynamics

**DOI:** 10.1101/583187

**Authors:** Jonas Windhager, Amelia Paine, Patroula Nathanailidou, Eve Tasiudi, María Rodríguez Martínez, John Lygeros, Zoi Lygerou, Maria Anna Rapsomaniki

## Abstract

DNA replication, the basis of genetic information maintenance, is a remarkably robust yet highly stochastic process. We present a computational model that incorporates experimental genome structures and protein mobility dynamics to mechanistically describe the stochastic foundations of DNA replication. Analysis of about 300,000 *in silico* profiles for fission yeast indicates that the number of firing factors is rate-limiting and dominates completion time. Incorporating probabilistic activation and binding, a full-genome duplication was achieved with at least 300 firing factors, with the only assumption that factors get recycled upon replication fork collision. Spatial patterns of replication timing were reproduced only when firing factors were explicitly activated proximally to the spindle pole body. Independent *in vivo* experiments corroborate that the spindle pole body acts as a replication activator, driving origin firing. Our model provides a framework to realistically simulate full-genome DNA replication and investigate the effects of nuclear architecture.

## Introduction

DNA replication is a tightly regulated process that ensures the faithful duplication of the genome before cell division. In eukaryotes, DNA replication initiates from multiple sites along the genome called *replication origins*. During G1 phase, the origin recognition complex (ORC) binds to replication origins and recruits the loading factors Cdt1 and Cdc6 [1, 2], and the six-subunit replicative helicase MCM2-7 [3, 4]. Together, these molecules form the pre-replicative complex (pre-RC) in a process known as *origin licensing* [5, 6]. *After the G1-S transition, a cascade of phosphorylation events, orchestrated by cyclin-dependent kinase (CDK) and Dbf4-dependent kinase (DDK) [7], leads to the binding of additional replication initiation factors* (e.g., Cdc45, Sld2, Sld3 and the four-subunit GINS complex [8, 9]) to the pre-RCs of selected origins, forming the pre-initiation complex (pre-IC) [10–12]. These events trigger *origin firing* [13], concluding with the formation of two sister replisomes (*replication forks*) that move in opposite directions, simultaneously unwinding the DNA and synthesizing a new copy [14, 15]. It has been recently suggested that firing factor complexes form dimers, which, upon origin firing, monomerize and travel along with each replication fork, and will re-dimerize when released following fork conversion [16, 17].

DNA replication in eukaryotic cells is a complex and uncertain process. Origin firing is stochastic and replication forks must progress continuously and replicate every part of the genome once and only once. In every cell cycle, a subset of all putative origins fire asynchronously, while the rest remain dormant until passive replication [18–21]. A variety of factors have been shown to influence DNA replication timing [22–24]. In fission yeast, replication origins are located in intergenic, nucleosome-depleted regions with high AT content [25–29]. Chromosomal context and local chromatin state are major determinants, with euchromatic and pericentromeric regions typically containing early-firing origins, and heterochromatic and subtelomeric regions containing late-firing and/or dormant origins [30, 31]. As recently observed in various organisms through chromosome conformation capture techniques, DNA replication timing is highly correlated with chromatin folding and global nuclear architecture [31–38]. In fission yeast, a gradient of origin efficiency centered at the vicinity of the spindle pole body (SPB) has been proposed, consistent with a diffusion-based mechanism coordinating DNA replication [38, 39].

A number of mathematical and computational models have attempted to explain the complex nature of DNA replication using different mathematical formulations, such as the KJMA framework [40, 41], anomalous reaction-diffusion and inhomogeneous replication kinetics [42, 43], probabilistic models of origin firing [44–46] and stochastic hybrid models [47, 48]. Recently, three-dimensional (3D) models have also been proposed. Using random origin locations in HeLa cells and random loop folding of chromatin, Löb et al. [49] were able to reproduce experimentally observed patterns in 3D. Using a spatiotemporal model, Arbona et al. [50] showed that the diffusion of firing factors alone is sufficient to account for characteristic replication kinetics captured by the probability of firing an origin per unit time and length of unreplicated DNA *I*(*t*), a function that forms the basis of many mathematical models of DNA replication [40, 51, 52]. However, none of the existing models have explored the effects of nuclear architecture on DNA replication dynamics using experimentally-derived genome structures, and the exact mechanisms governing replication initiation and progression remain unclear.

In this work we set out to investigate the interplay between linear DNA replication, protein mobility dynamics and spatial chromatin structure. We present an integrated model for DNA replication that exploits 3D genome structure and 3D origin coordinates as derived by chromatin conformation capture experiments to simulate the spatiotemporal dynamics underlying origin firing. By explicitly modeling nuclear protein mobility, probabilistic origin firing and linear replication fork progression, we incorporate diffusion and binding dynamics to mechanistically describe the stochastic foundations of DNA replication. Analysis of a large number of *in silico* replication profiles enables the assessment of model parameter sensitivity and reveals spatial patterns that were independently assessed *in vivo*. In summary, our model provides a framework to realistically simulate full genome DNA replication, and to investigate the relationship between 3D chromatin conformation and DNA replication timing.

## Results

### A stochastic 3D model of DNA replication

We present a spatiotemporal model of DNA replication that incorporates both protein and replisome movement within the nucleus to capture the stochastic nature of DNA replication timing. The model consists of two interlinked modules: a **particle-centric** module that models the mobility of initiation factors within the nucleus and an **origin-centric** module that captures origin firing and linear progression of replisomes along the genome. Both modules are characterized by hybrid dynamics (e.g., continuous movement of replication forks, discrete binding/unbinding events) and stochasticity (e.g., firing factor diffusion, origin firing). Our model does not require experimentally-derived origin firing probabilities as input since it explicitly models origin firing as the interaction between initiation factors and origin sites. The spatiotemporal model simulates DNA replication at a whole-genome level and is tailored to the case of fission yeast, exploiting recent 3D genome structures derived by chromosome conformation capture experiments [39, 53]. Due to its conserved features, fission yeast has long served as an attractive model organism for the study of DNA replication. Furthermore, its relatively small genome of approximately 14 megabases and three chromosomes allows for whole-genome simulations within reasonable time.

A graphical summary of the model is given in Fig. 1, and a full mathematical description is given in Methods. A number of experimentally derived input parameters are required (Fig. 1A), with which the model can be extended to any eukaryotic organism. Model simulations yield *in silico* DNA replication profiles and allow the exploration of DNA replication dynamics for different hypotheses and parameter values. An animation of an example simulation can be found in the Supplementary video.

**Fig. 1:**
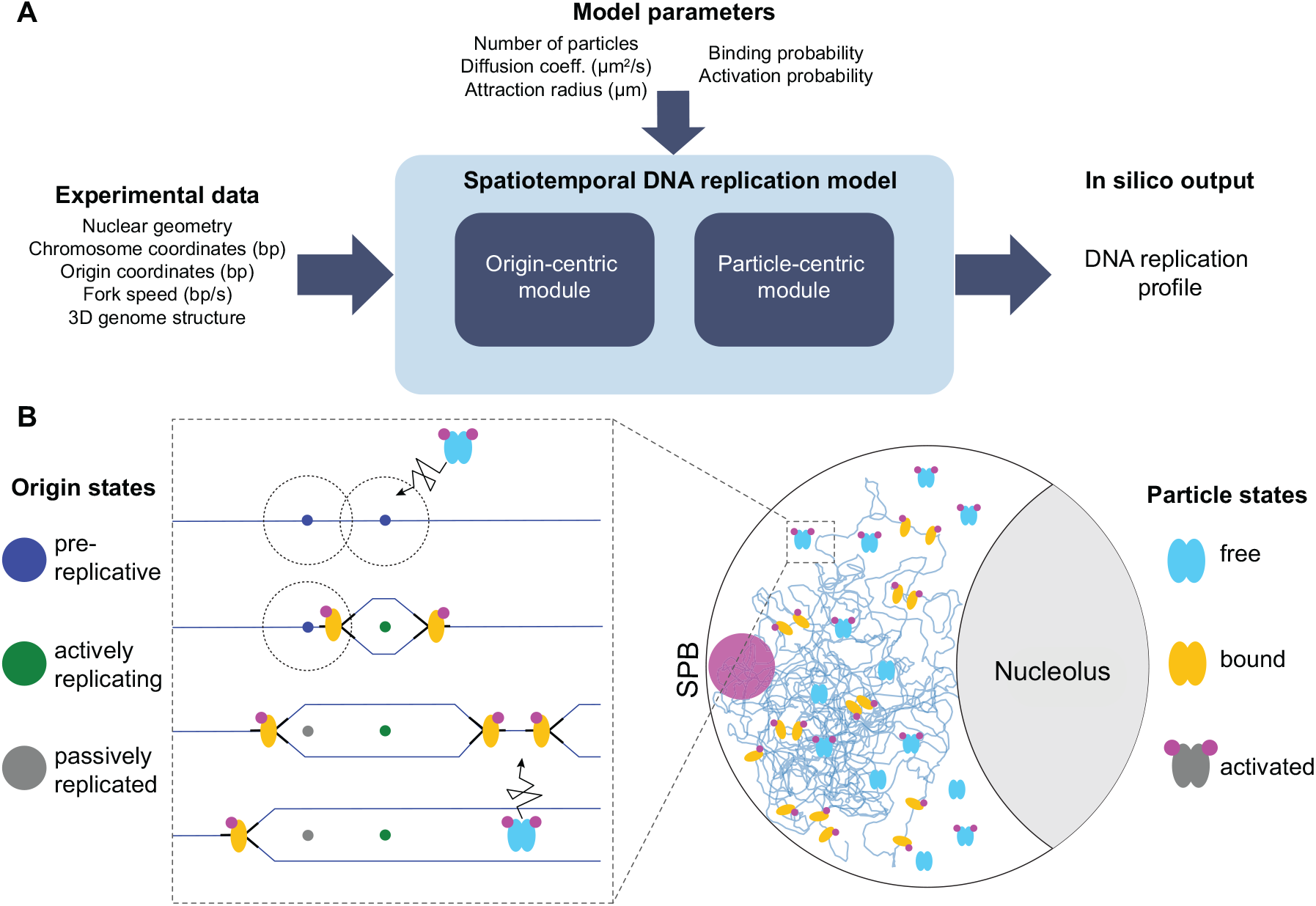
(A) Overview of the DNA replication model. Simulations with the required inputs give rise to *in silico* DNA replication timing profiles. **(B) Schematic representation of the two modules**. Right: Firing factor particles found at different discrete states diffuse within the nucleus and activated particles bind to origins. Left: An unbound and activated particle enters the attraction sphere (dotted circle) of a licensed replication origin, initiating firing and changing the discrete origin state to actively replicating. Two replication forks emanate from the origin and move along the linear DNA sequence in opposite directions, replicating the genome. The particle splits into two monomers that travel along with the replication forks. Licensed origins that have not fired before a fork passes through switch state to passively replicated. When forks moving in opposite directions collide, they merge and the particle dimers are released to freely diffuse and potentially fire other licensed origins.

### Nuclear architecture

Following [39, 53], the fission yeast nucleus is modeled as a 3D spherical domain with impermeable boundaries excluding the nucleolus (Fig. 1B – right and Methods). The spindle-pole body (SPB) is localized diametrically opposite of the nucleolus and anchors centromeres in its vicinity, forming a specialized nuclear subdomain [54, 55]. Spatial origin coordinates were estimated by mapping the experimentally determined base-pair coordinates of Heichinger et al. [29] onto the 3D whole-genome coordinates of Grand et al. [53] and Pichugina et al. [39].

### Particle-centric module

The particle-centric module of our model represents the behavior of the initiation factors (hereafter referred to as particles) within the nucleus. For simplicity, we model the cascade of molecular interactions in origin firing as a binding event by a single collective particle. Each particle is characterized by three states: a continuous state representing the position of the particle in the nucleus and two discrete binary states representing the particle’s activation and binding states, respectively (Fig. 1B – right and Supplementary Fig. S1A). Particles are assumed to already be dimers and diffuse freely in the nucleus if not bound to an origin. Similar to [56, 57], particle diffusion is described by a stochastic differential equation and assumed to be isotropic, space-homogeneous, and effective. The latter implicitly accounts for biophysical interactions, such as collisions with macromolecules or transient binding to DNA. A particle switches from the inactive state to the active state through phosphorylation when passing at the vicinity of the SPB, and between the free and bound states through its interaction with origins.

### Origin-centric module

The origin-centric module of our model is largely based on the stochastic hybrid model of Lygeros et al. [47]. Briefly, a replication origin can be in one of the following discrete states at any time: *Pre, RB, RL, RR, Post*, or *Pass* (Supplementary Fig. S1B). Initially, all origins are assumed to be licensed and in the *Pre* (pre-replicative) state. Upon firing, replication forks start traveling in either direction, and the origin enters the *RB* state (replicating bidirectionally). When a fork collides with a fork emanating from another origin moving in the opposite direction, the initiating origin changes state from *RB* to *RL* or *RR* (replicating to the left / right) depending on whether the left or right fork is still active. Once both forks have collided and stopped replicating, the initiating origin switches to the *Post* (post-replicative) state and does not contribute to the replication process anymore. If a replication fork passes through a *Pre*-origin that has not fired yet, the origin is blocked from future firing to prevent re-replication of DNA and enters the *Pass* (passively replicated) state. Replication is complete when all origins are in the *Pass* or *Post* state, which implies that all forks have either merged or reached the end of a chromosome.

### Particle-origin interaction

The crosstalk between the two modules is orchestrated by particle-origin interactions (Fig. 1B – left). For origin firing to occur, an unbound and activated particle must diffuse into the attraction sphere of a *Pre* state-origin and bind to it (Supplementary Fig. S1C). To capture the inherent stochasticity of this process, a parameter *P*_*bind*_ is introduced, representing the probability of a particle binding to an origin when it is within range. Upon firing, the particle splits into two monomers that travel along the genome with the replisomes. When two replication forks converge, the reassembled dimer is released to freely diffuse in the nucleus and potentially fire other origins (Supplementary Fig. S1D). Particle state transitions are governed by stochastic dynamics, apart from the bound →free transition, which occurs when forks merge and is thus deterministic given the time of firing. Origin state transitions are governed by deterministic dynamics, apart from the *Pre*→ *RB* transition, which is governed by the stochastic dynamics of particle binding. The tight interplay between the two modules is thus executed through these interlinked transitions, with particle dynamics affecting origin dynamics and *vice versa*.

## Model variants

To examine how nuclear architecture affects the replication process, we conducted multiple simulations under the following conditions:

1. **Random origin positioning:** The model is instantiated with genome structures generated randomly; structures are simply confined in the nucleus, but not fitted to experimental data [53, “confined” model]. Particles are initialized uniformly in the nucleus in the active state.
2. **Realistic origin positioning:** To examine whether origin localization affects replication timing, 3D genome structures generated from genome conformation capture experiments are used as input [53, “interactions” model], selected as described in Methods. Particles are initialized as in model variant 1.
3. **SPB-mediated particle activation:** Experiments have shown that SPB-proximal origins fire earlier, possibly due to the activation of initiation factors in the SPB-proximal nuclear region, followed by firing of origins encountered as they diffuse [39]. To model this process, particles are initialized in the inactive state and become activated within the SPB-proximal region. To account for the stochastic nature of the underlying biochemical process, for each diffusion step within the SPB-proximal region, there is a constant probability of activation (*P*_*act*_) if the particle has not yet been activated. SPB-mediated particle activation slowly introduces activated particles to the nucleus and is equivalent to modeling an influx of particles from outside the nucleus through the SPB. Experimentally-derived genome structures are used as input, same as in model variant 2.

### Implementation and simulations

The numerical implementation and simulation of the continuous model was executed by gridding time and space [56] (Methods). The joint model was implemented in a highly parallelized and modular fashion using C++ and MPI, with optimizations to further increase simulation throughput. Simulation visualizations and source code are available at https://github.com/AI4SCR/dna-replication.

### Analysis of *in silico* replication profiles

An initial set of Monte Carlo simulations was conducted for all model variants and a broad range of parameter values (Supplementary Table S1). Parameter values for the origin attraction radius *r*_*attr*_ were estimated based on crystal structures of the origin recognition complex [58] and the MCM2-7 double hexamer [59] that, when arranged next to each other, form a ≈ 0.008 *μm*-long complex. Under the simplifying assumption that activation factors can dock anywhere on the surface of the two pre-RCs bound to each origin, we set the minimal value of the origin attraction sphere radius to 0.015 *μm* and explore higher values as well. To select plausible values of the particle diffusion coefficient, we considered that in the fission yeast nucleus the diffusion coefficient of GFP alone was reported to have a minimum value of ≈ 3*μm*^2^*/s* [60], and sampled three realistic values in the range of [0, 3.5]*μm*^2^*/s*. The particle activation and binding probabilities were set to *P*_*act*_ = *P*_*bind*_ = 1. To account for the stochastic nature of the replication process, five iterations were performed for each structure and parameter combination, resulting in a total of 192,240 individual simulations. For each simulation, replication timing statistics were recorded.

#### DNA replication completion time

The experimentally established time until completion of DNA replication in fission yeast [61] is an important indicator of the model’s ability to reflect biological reality. The results of the initial simulations show that the completion time is largely dominated by the number of particles present in the nucleus, with completion time decreasing rapidly as *N* increases (Fig. 2A). Notably, at high *N*, the completion time converges to the minimum theoretical completion time of 652.92 seconds, equivalent to the time needed to replicate the maximum inter-origin distance [29]. Simulations with *N* = 150 achieved the reported completion time of 20 minutes in fission yeast [61]. The effect of the diffusion coefficient and the attraction radius on the distribution of completion times was negligible when *N* was held constant (Fig. 2B, 2C), despite the exploration of a broad range of plausible values for both parameters. Similarly, the model variant did not have a noticeable impact on the completion time. Overall, the timescale of particle diffusion throughout the nucleus is much shorter than that of replication forks moving along DNA strands.

**Fig. 2:**
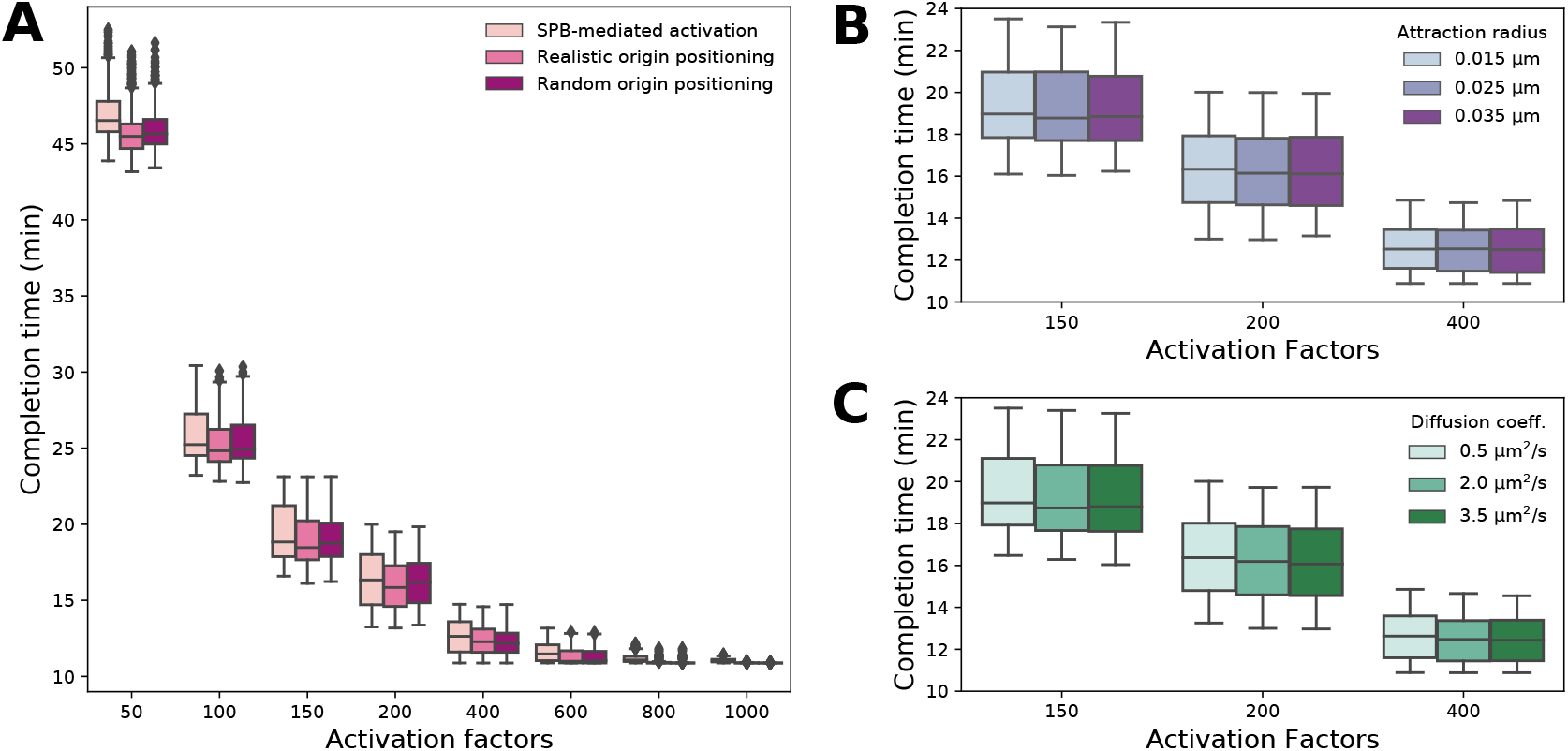
Completion time distributions for different model variants and simulation parameters. Simulated completion times across a wide range of number of particles *N* for the three model variants **(A)**, different attraction radii *r*_*attr*_ **(B)** and different effective diffusion coefficients *D* **(C)**. Unless otherwise specified, the SPB-mediated activation variant is used, *D* = 0.5*μm*^2^*s*^*−*1^, and *r*_*attr*_ = 0.025*μm*. Relevant probabilities are set to 1.

#### Spatial organization of origin firing

To examine the spatial organization of replication in the nucleus, relative radial kernel density estimates (KDEs) were calculated for experimentally-derived [29] and simulated origin firing times (Fig. 3, Methods). The random origin positioning model variant shows a uniform, random distribution of early and late-firing origins, with no discernible pattern. The same is true for the realistic origin positioning variant; genome structure alone is not sufficient to reproduce the experimental data, suggesting that an additional mechanism is in place. Indeed, the SPB-mediated particle activation variant shows a similar firing pattern to the experimental data, with early-firing origins located close to the SPB and late-firing origins close to the periphery and nucleolus. Due to simplifying assumptions such as equal binding probabilities for all origins and single binding events to fire origins, the early firing effect appears stronger in simulations than in experiments, to the point of depleting intermediate origins in the SPB region. Alternative types of KDE plots reveal efficient yet late-firing origins near the nucleolus in addition to those near the SPB, suggesting a wave-like origin firing behavior (Methods, Supplementary Fig.S3).

**Fig. 3:**
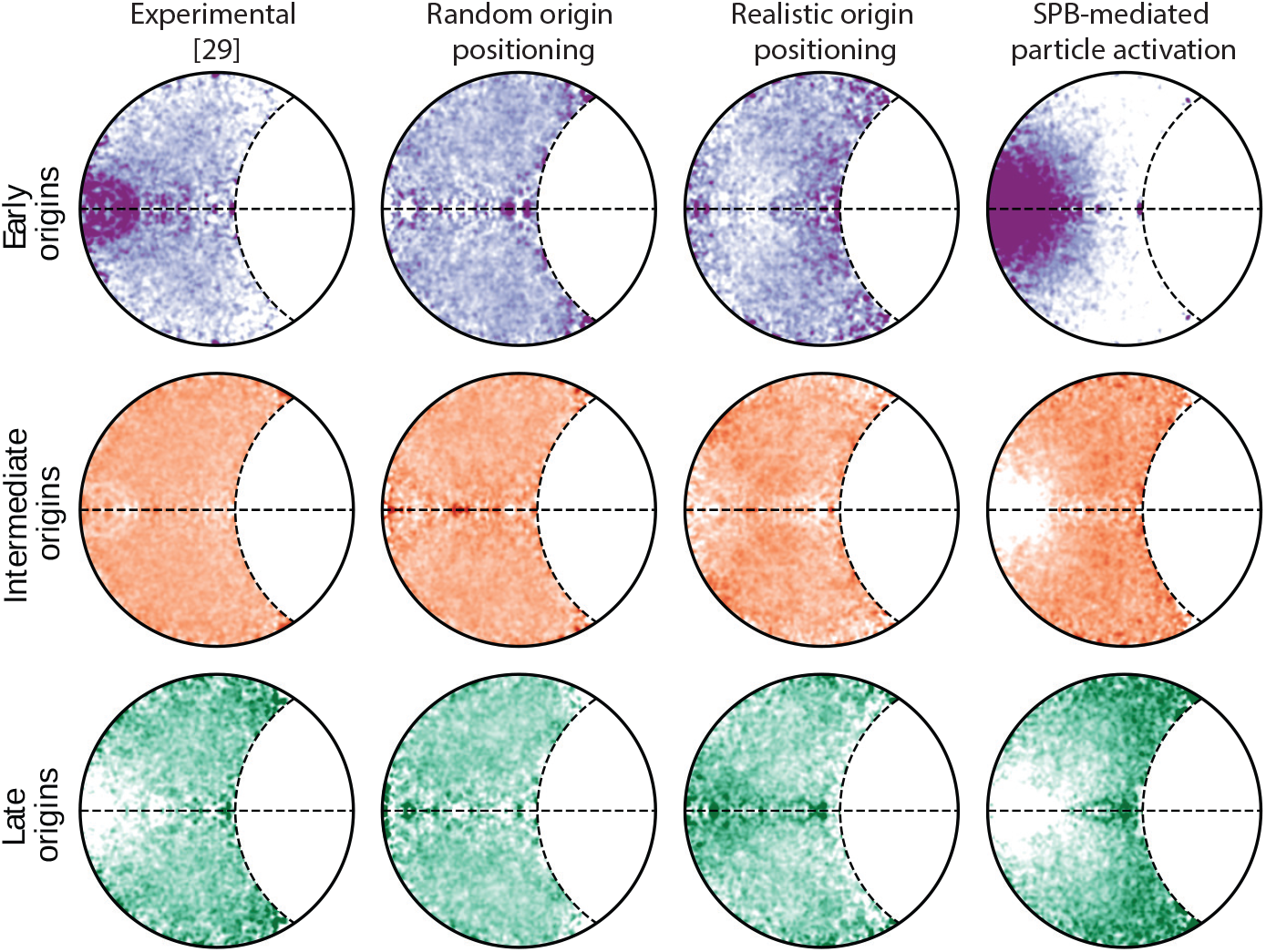
Spatial organization of average origin firing times. Relative radial kernel density estimates of earlyfiring (top row), intermediate (middle row) and late-firing (bottom row) origins for experimental data and all three model variants. To characterize origins, we ranked them by their firing time and assigned origins that belonged to the lower quartile (Q1) as early, origins with *Q*_1_ *<* firing time *< Q*_3_ as intermediate, and origins in the higher quartile as late.

#### Exploration of model probabilities

As the number of particles *N* has been shown to be an influential model parameter and the SPB-mediated particle activation best reproduces the experimental patterns of origin firing, a second set of simulations of this model variant was conducted to explore *N* in more detail and observe the effect of different values of the activation and binding probabilities. Model inputs were set as in Supplementary Table S1. We first examined the effect of different model probabilities on the DNA replication completion times for various values of the number of particles *N*, shown as heatmaps in Fig. 4. Many combinations achieved the desired completion time of 20 minutes, generally showing a trade-off between activation probability *P*_*act*_ and binding probability *P*_*bind*_ to maintain the same completion time. As expected, decreasing *P*_*act*_ or *P*_*bind*_ increases the overall completion time, so the completion time is highly tunable for any value of *N*.

**Fig. 4:**
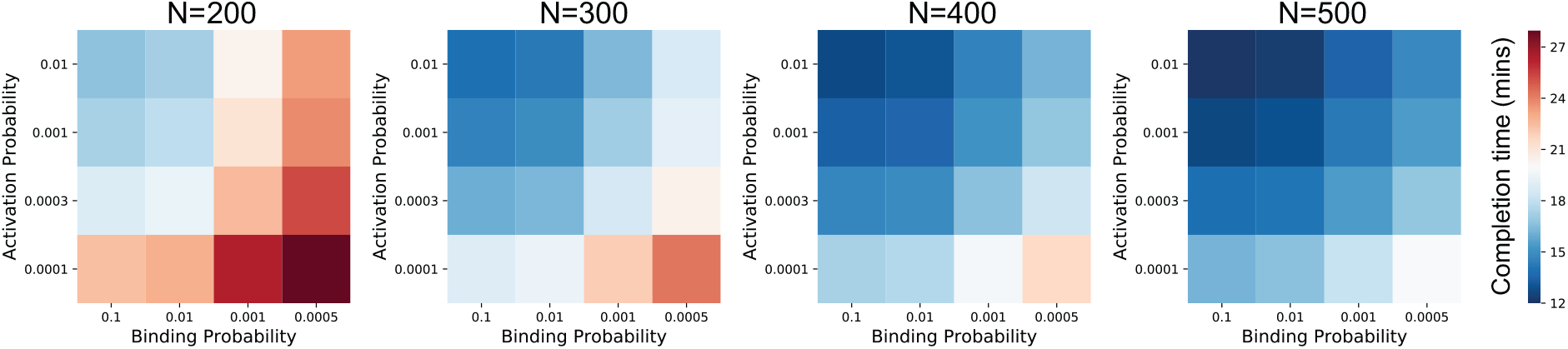
Sensitivity analysis of model probabilities. Mean completion time (in minutes) for various combinations of model parameters and numbers of particles *N*, shown on a diverging color scale centered on the desired completion time of 20 minutes.

#### Origin firing concurrence

Recently, experimental evidence for the presence of clusters of fired origins in late S phase, organized in nuclear replication foci, has been reported [62]. To assess this effect *in silico*, we examined the tendency of all origin pairs to fire concurrently: for each pair of origins, we computed a firing concurrence score, where high values indicate that only one origin of the pair fires at a time, and low values indicate that they exhibit the same behavior (they either both fire or do not fire at all) (Methods). As shown in Fig. 5 for Chromosome 3, most origin pairs are uncorrelated, but patterns emerge among origins in linear DNA proximity. When *P*_*bind*_ is relatively large (Fig. 5A – left), the blue halo around the diagonal of the concurrence matrix indicates that neighboring origins are likely to fire together, since particles initially tend to bind to SPB-proximal origins. Box plots of firing concurrence indicate that origin firing is correlated within a neighborhood of about five origins (Fig. 5B). Immediate neighbors exhibit a competitive relationship, obvious from the red super- and sub-diagonals of the concurrence matrix, as when one origin fires it passively replicates its nearest neighbors and prohibits them from firing. Conversely, when *P*_*bind*_ decreases (Fig. 5A – right), the positive effects are replaced by strong competition within an origin’s neighborhood, indicated by both strong red around the diagonal and low firing concurrence values up to the three nearest neighboring origins: particles tend to diffuse throughout the nucleus instead of firing adjacent origins, so fired origins can passively replicate more neighbors before fork collision.

**Fig. 5:**
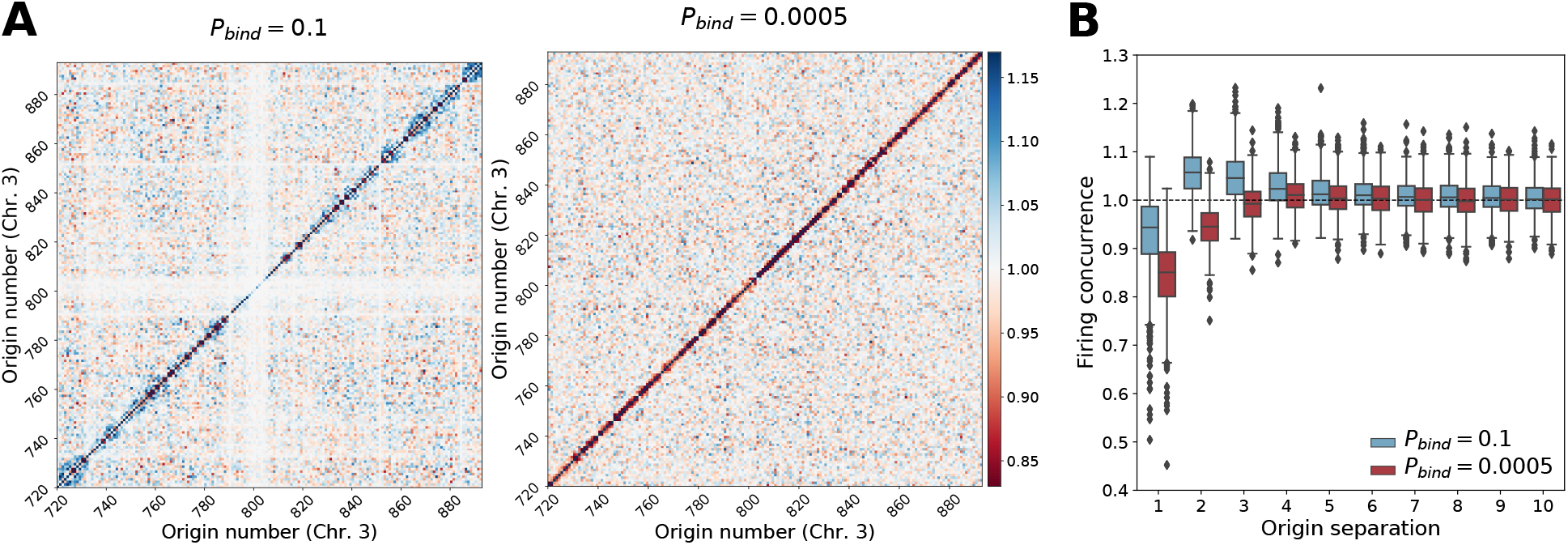
Neighborhood effects on origin firing. In all parameter sets, *N* = 150, *D* = 0.5*μm*^2^*s*^*−*1^, *r*_*attr*_ = 0.025*μm*, and *P*_*act*_ = 0.01. **(A)** Concurrence matrices of Chromosome 3 for high and low values of *P*_*bind*_. **(B)** Distributions of firing concurrence for neighboring origins, demonstrating local concurrence when *P*_*bind*_ = 0.1 and strong local competition when *P*_*bind*_ = 0.0005.

### *In vivo* exploration of spatial organization

To monitor the spatial distribution of early replication *in vivo* at the single-cell level, we performed immunofluorescence against the thymidine analog IdU, which marks nascent DNA, in fission yeast cells arrested in early S phase using hydroxyurea (Methods). Under these conditions, efficient origins fire [29] and will incorporate IdU. Cells were counter-stained with Hoechst, which labels all nuclear DNA. The strain used also expresses the centromeric protein Mis6 tagged with GFP (Mis-GFP). In normal cells, centromeres are clustered near the nuclear periphery at the site of the SPB and a single Mis6-GFP dot is observed [63]. As shown in Fig. 6A, IdU incorporation is restricted to nuclear areas surrounding Mis-GFP. This is consistent with origins firing at the proximity of the SPB-clustered centromeres in early S-phase. We used a mutant strain (Csi1*Δ*) to uncluster centromeres and disassemble then from the SPB [64]. In Csi1*Δ* cells, IdU incorporation appears much more dispersed through the nucleus, and the IdU/Hoechst area ratio increases significantly (p-value = 1.5 · 10^*−*4^) (Fig. 6A, B). This is consistent with SPB-clustered centromeres acting as a positive pole for replication initiation. To more thoroughly investigate the correlation between distance from SPB and origin efficiency, we tagged one efficient and one inefficient origin [29] in each of the three fission yeast chromosomes, using an array of lacO repeats inserted in the genome (Fig. 6C, Methods). These strains also expressed lacI tagged with GFP, which binds to lacO repeats and marks the location of the lacO-tagged origin, and Mis6-mCherry, which marks the SPB-clustered centromeres. The 3D Euclidean distance between the SPB and each tagged origin was measured microscopically in hundreds of cells (Fig. 6D). In fission yeast cohesin-mediated genome folding in small domains (75-100 Kb) is retained throughout interphase [65], while condensin-mediated long-range interactions (0.3-1 Mb) are established by mitotically expressed transcription factors and therefore are specific to mitosis [65–67]. These studies show that major genome reorganization happens only during mitosis, which represents a small fraction (10%) of the cell cycle, and that genome structure during G1, S and G2 is subjected to subtle changes. Given this we reasoned that asynchronous populations of cells would reflect the genomic organization during S phase and thus distances were measured in asynchronous cells. Efficient origins (orange) are closer to the SPB than inefficient ones (blue) (p-values = 5.6 · 10^*−*6^, 5.3 · 10^*−*7^, and 6.3 · 10^*−*8^ for origin pairs in Chromosomes 1, 2, and 3, respectively). The efficiency of each origin was independently verified by real-time quantitative PCR (qPCR, Fig. 6E). Our analysis in fission yeast cells is in agreement with the simulations of the SPB-mediated particle activation model (Supplementary Fig. S4), and supports the SPB as a source of a replication activator which drives origin firing.

**Fig. 6:**
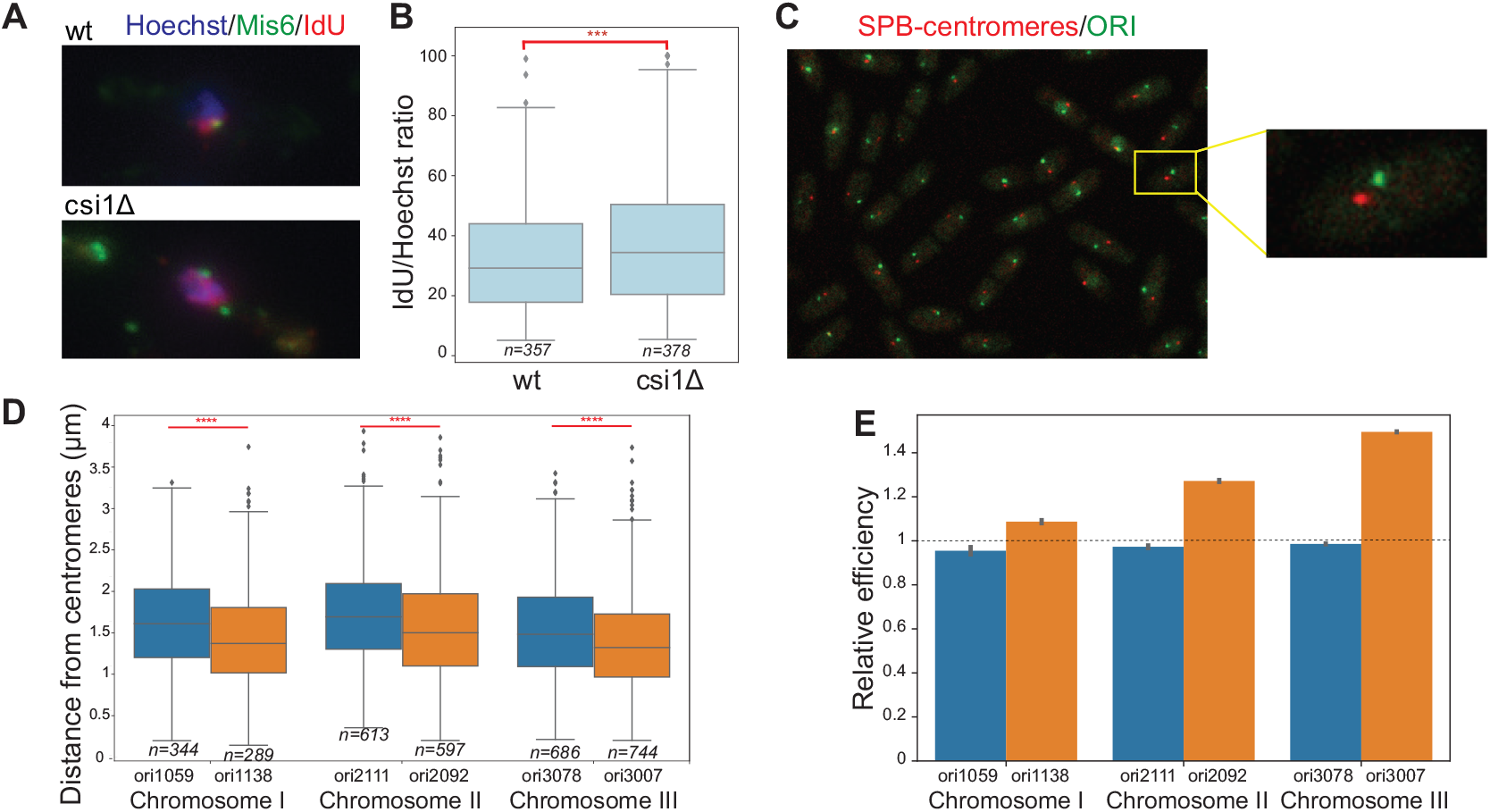
*In vivo* exploration of spatial organization. **(A)** Immunofluorescence against IdU in Mis6-GFP wild type and Mis6-GFP Csi1*Δ* fission yeast cells arrested in early S using HU. Mis6 marks the centromeres (green), IdU stains nascent DNA (red) and Hoechst stains the whole nucleus (blue). **(B)** IdU/Hoechst area ratio (in percent), plotted for wild type and Csi1*Δ* cells. **(C)** Tagging of specific origins using the lacO-lacI system in a Mis6-mCherry background strain. A representative image is shown. **(D)** For each chromosome, the 3D distance to the centromeres/SPB structure for one efficient (orange) and one inefficient (blue) origin was measured microscopically. **(E)** The efficiencies of the respective origins were validated by real-time qPCR. All p-values were computed using a standard Mann-Whitney statistical test.

### DNA replication kinetics

We next assessed how model probabilities affect DNA replication kinetics in terms of mean replication rate (Fig. 7A, Methods). High values of *P*_*act*_ and *P*_*bind*_ lead to almost all particles activating and binding within the first minutes and quickly binding new origins upon release, resulting in a relatively constant and high replication rate that abruptly decreases shortly before replication completes. Binding probability primarily affects mid to late S phase. Low values of *P*_*bind*_ delay the particles from binding to origins and increase the pool of free particles, resulting in a lower maximum replication rate reached and a more gradual decline. Conversely, decreasing *P*_*act*_ has a prominent effect on early S phase, as it delays and decreases the maximum replication rate.

**Fig. 7:**
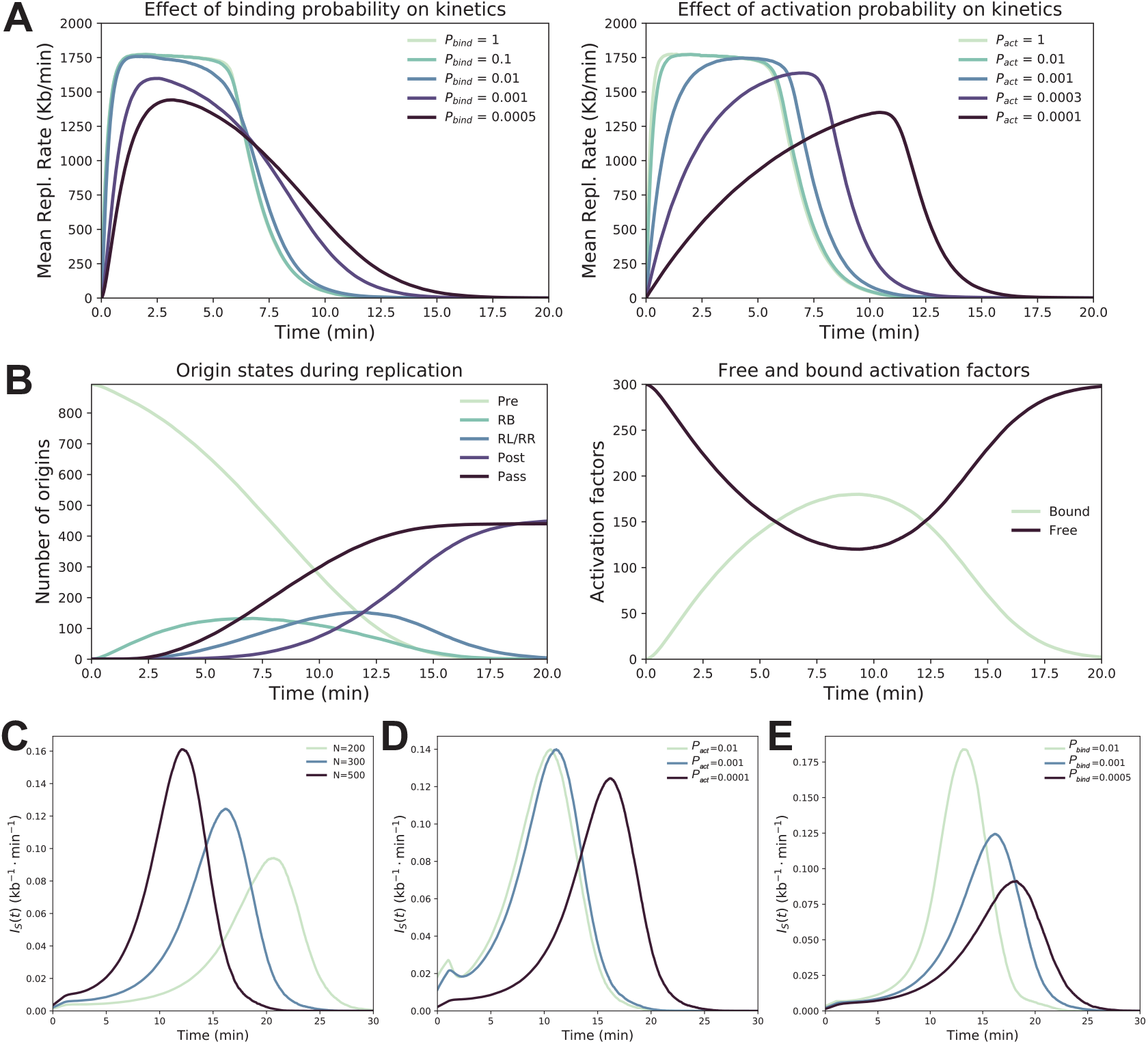
Summary of replication kinetics for different parameter values. In all simulations, *N* = 300, *r*_*attr*_ = 0.025*μm*, and *D* = 0.5*μm*^2^*s*^*−*1^. **(A)** Effect of *P*_*bind*_ (*P*_*act*_ = 0.01) and *P*_*act*_ (*P*_*bind*_ = 0.1) on mean replication rate. **(B)** Average number of origins in each model state over the course of replication for a single parameter set (*P*_*bind*_ = 0.001, *P*_*act*_ = 0.0001) and the corresponding average number of free and bound particles. **Simulated firing rate per length of unreplicated DNA** *I*_*S*_ (*t*). Mean *I*_*S*_ (*t*) for a variety of parameter sets (varying *N* **(C)**, *P*_*act*_ **(D)** and *P*_*bind*_ **(E)**), resembling the characteristic bell shape observed in eukaryotes [68]. In all cases, *r*_*attr*_ = 0.025*μm, D* = 0.5*μm*^2^*s*^*−*1^. *N* = 300, *P*_*bind*_ = 0.001, and *P*_*act*_ = 0.0001, unless otherwise specified.

Tracking the number of origins and particles in each state upon probabilistic activation and binding provides insight into replication kinetics (Fig. 7B). The number of origins in the *Pre* state steadily decreases from 893 to 0 throughout replication. The number of origins in the *RB* state peaks before *RL* or *RR*, as expected, because forks begin to collide a few minutes after origins start firing. Most passive replication occurs during mid S phase, whereas the transition to *Post* occurs primarily during mid to late S phase, when the replication rate is decreasing. The number of bound particles, proportional to the replication rate, exhibits a bell-shaped profile with a peak in mid S phase, agreeing with experimental evidence [62].

The “probability of initiating an origin per unit time and per length of unreplicated DNA” introduced by Herrick et al. [68] forms the basis of many mathematical models of DNA replication [40, 51, 52]. This function, denoted *I*(*t*), where *t* is the time since the start of S phase, resembles a bell-curve shape for eukaryotes, peaking in mid-to late-S phase. Goldar et al. [40] developed a mathematical model postulating the presence of a limiting factor that binds to unreplicated locations of the DNA initiating replication and is released upon fork convergence. Arbona et al. [50] demonstrated that a bell-shaped *I*(*t*) arises from a DNA replication model incorporating a fixed number of firing factors that bind to origins, move along the DNA, and release upon fork collision. We reproduced the characteristics of *I*(*t*) using the “simulated firing rate per length of unreplicated DNA” *I*_*S*_(*t*) described by Arbona et al., which is proportional to *I*(*t*) [50] (Methods). Note that the simulated firing rate should be proportional to the number of free activated particles and the number of pre-replicative origins, which helps to explain the overall behavior of *I*_*S*_(*t*). The results for a variety of parameter sets are shown in Fig. 7D-E. The small increase of *I*_*S*_(*t*) in early S phase is accounted for by particle activation, since both the number of *Pre*-origins and amount of unreplicated DNA do not change appreciably relative to their magnitudes. After mid S phase, the amount of unreplicated DNA diminishes rapidly, resulting in a sharp increase and peak of *I*_*S*_(*t*). At the end of replication, the number of *Pre*-origins approaches zero, causing a rapid decline of *I*_*S*_(*t*). Increasing *N* or decreasing *P*_*bind*_ delays the peak and reduces its maximum height. Notably, high values of *P*_*act*_ generate a small peak at the beginning of replication, because the immediate presence of many activation factors at the start results in a spike of initial firing activity. The overall behavior of *I*_*S*_(*t*) is not sensitive to parameter changes within the parameter ranges we explored, confirming that a bell-shaped *I*(*t*) curve arises from a limited number of particles in conjunction with recycling of firing factors upon fork collision.

## Discussion

In this work a spatiotemporal model of DNA replication is presented. The model simulates the diffusion of firing factor particles within the nucleus, their probabilistic interaction with replication origins that leads to origin firing, and the movement of replication forks along the genome, capturing the interlinked stochastic hybrid dynamics that govern DNA replication. Although an exhaustive search for the best-fitting parameter set was out of scope, our simulations provide insight on plausible parameter ranges.

The number of particles *N* strongly affected the time needed for a full DNA replication, whereas the origin attraction radius and the diffusion coefficient had negligible effects. The latter is likely attributed to different timescales of diffusion and linear DNA replication: even with a low diffusion coefficient of *D* = 0.5*μm*^2^*s*^*−*1^, a particle would traverse the 2.66*μm* nuclear diameter in about seven seconds, a much smaller timescale than that of the replication process. For instantaneous particle activation and binding, 150 particles were enough to complete replication within the experimentally established 20 minutes limit [61]. However, the underlying kinetics exhibited patterns markedly different to experimental observations. Incorporating probabilistic particle activation and binding extended the time to replication completion, but increasing the number of particles counteracted this effect. Indeed, for lower values of *P*_*act*_ and *P*_*bind*_, a full genome duplication was achieved with at least 300 particles, consistent with up to 400 replicons predicted through DNA combing experiments, and the recorded replication rate over time matched experimental observations [62]. Additionally, our model reproduced the characteristic probability of initiating an origin per unit time and length of unreplicated DNA with the only assumption that the number of particles is rate-limiting. This is in agreement with extensive experimental evidence from the literature, reporting that factors participating in origin firing, such as DDK, Sld2, Sld3 and Cdc45, exist in limiting quantities [22, 69–72]. It is also consistent with our prior *in silico* analysis [47], where the presence of a limiting factor that is continuously redistributed in origins during replication is a plausible explanation for the so-called random gap problem [73].

Our model was able to capture global patterns of origin firing, manifested as a gradient of origin efficiency emanating from the SPB [39], when particle activation at the vicinity of the SPB was modeled explicitly. Nuclear architecture proved to be essential through independent *in vivo* analysis, agreeing with SPB’s role as a positive pole in replication initiation, which was lost when centromere clustering was disrupted.

These findings indicate that genome structure alone is not sufficient to reproduce the observed patterns of origin firing, and that firing factors must diffuse to the SPB-proximal region to get activated possibly through their phosphorylation by different kinases before they can bind to origins. Indeed, evidence for the centromeric localization of DDK has been reported for both yeast species. In fission yeast, DDK is recruited to pericentromeres by the epigenetic regulator Swi6, and in budding yeast by the Ctf19 kinetochore complex [74–77]. Alternatively, firing factors could be introduced in the nucleus already in the active state through SPB-specific nuclear pores, but, to our knowledge, nuclear pores have not previously been reported to have such a role.

Conversely, more fine, local patterns of origin firing, such as experimentally observed replication foci [62], could not be reproduced, as in our simulations firing concurrence was observed for larger values of the binding probability only between linearly-proximal origins. One possible explanation for this discrepancy lies in the genome structures themselves: they are derived from a single, population-level chromosome conformation map, which lacks the resolution that would allow capturing local chromatin folding in fine detail, and cannot faithfully represent the underlying cell-to-cell heterogeneity. Similarly, the origin locations used as input to our model were experimentally determined by pooling a cell population and thus might again mask the underlying heterogeneity, implying that only the most frequent origin sites might be detected, or that origin clusters might appear as a single origin [78, 79]. As single molecule methods mature and become more widespread [80, 81], the emerging data can lead to models with more accurate predictions.

In our work we explored how chromatin structure affects replication timing and pointed to a positive correlation between SPB proximity and early firing. However, origin efficiency has long been correlated with sequence characteristics, such as AT content or intergenic size [25–29, 82, 83]. It still remains to be seen if a causal link between sequence and structure exists, or if they are two independent mechanisms of origin firing regulation. At the same time, the constant, uniform binding probability used in our model may be too simplistic to capture local patterns of DNA replication, and more complicated mechanisms could be in place. Origins may start with non-uniform binding probabilities, scaled based on sequence characteristics, which may dynamically evolve depending on replication progression or local chromatin accessibility. Simulating multiple particle species that diffuse within the nucleus, form multi-protein complexes, and progressively bind to and fire origins may be a more realistic scenario that implicitly reduces each origin’s firing efficiency depending on its location. Last, in our model we assume that particles diffuse already as dimers, but alternative hypotheses on the exact order of dimerization, activation and origin binding could be examined. Future extensions of the model will allow us to investigate these open questions.

## Acknowledgements

We thank Dr. Justin M. O’Sullivan and Dr. Tatyana Pichugina (Liggins Institute, University of Auckland, New Zealand) for their support in chromosome conformation capture data analysis and modeling. The work by J.L. was supported by SystemsX.ch under the project SignalX. We acknowledge support by the project Bioimaging-GR (MIS5002755) funded by the Operational Programme “Competitiveness, Entrepreneurship and Innovation” (NSRF 2014-2020) and co-financed by Greece and the European Union (European Regional Development Fund).

## Author contributions

Conceptualization and Supervision, J.L., Z.L., and M.A.R; Methodology, J.W., A.P., E.T., M.R.M, J.L. and M.A.R.; Software and Formal Analysis, J.W., E.T. and A.P.; Biological Experiments, P.N. and Z.L.; Investigation, J.W., A.P., P.N. and E.T.; Writing - Original Draft, J.W., A.P., E.T., P.N., and M.A.R.; Writing - Review and Editing, all authors.

## Methods

### DNA Replication model

The spatio-temporal DNA replication model is composed of two modules closely interacting with each other: A particle-centric module that simulates the mobility of firing factors within the nucleus, and an origin-centric module that simulates how replication origins change states and replication forks progress along the genome. All processes from both modules occur in parallel and take place within the nucleus.

In our work we tailored the nucleus to fission yeast based on a model by Grand *et al*. [53] and assumed it is in the S phase of the cell cycle. Briefly, the nucleus is modeled as a spherical domain 𝒩 with radius *r*_*nucleus*_ = 1.33*μm* and impermeable boundaries. The nucleus domain excludes a nucleolus region with a volume *V*_*nucleolus*_ relative to the nuclear volume *V*_*nucleus*_:

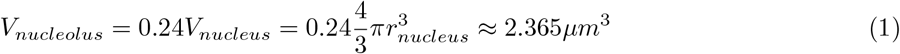

The nucleolus is modeled as the intersection of the nucleus with a displaced sphere of equivalent radius *r*_*nucleus*_. Given the height *h*_*cap*_ of the two spherical caps forming the intersection, the nucleolus displacement along the x-axis *Δx*_*nucleolus*_ is given by

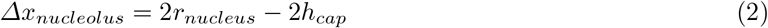

The spherical cap height *h*_*cap*_ is related to the *V*_*nucleolus*_ via the cap volume *V*_*cap*_:

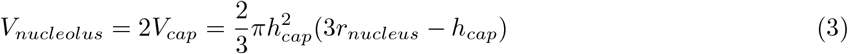

Using Cardano’s method, the resulting cubic equation can be solved for *h*_*cap*_, yielding a horizontal nucleolus displacement of *Δx*_*nucleolus*_ ≈ 1.510*μm*. With ∥**x**∥ being the Euclidean norm of **x**, the nucleus domain 𝒩 is thus given by:

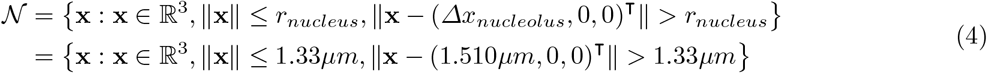

Furthermore, the nuclear domain𝒩contains a region proximal to the spindle pole body (SPB), which in fission yeast is the anchoring point for centromeres [55]. The SPB-proximal region is modeled as a sphere with radius *r*_*SP B*_ = 0.2*μm*, with a horizontal displacement of:

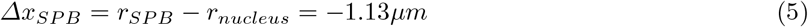

In the **particle-centric module**, the diffusion and binding of firing factors (*particles*) within the nucleus is simulated. Different types of particles are collectively represented by a single, generic type. The model is described by hybrid dynamics: continuous dynamics are associated with the movement of the particles within the nucleus, and discrete dynamics are associated with their binding and activation states. The evolution of the continuous and discrete states of the system are stochastic and interlinked: the continuous dynamics depend on the discrete dynamics and vice versa. Similar to stochastic hybrid models presented for other biological processes [56, 57], the particle-centric module follows the laws of a Jump Diffusion Process (JDP), where the continuous state evolves stochastically according to a stochastic differential equation (SDE) and the transitions between discrete states are modeled using probability distributions. A brief overview of the module is given in Supplementary Fig. S1A.

All particle copies are located within the nuclear domain 𝒩 and we assume that their count remains fixed throughout the process of replication. For each molecule *i* = 1, …, *N* at each time point *t*, let *p*_*i*_(*t*) = [*x*_*i*_(*t*) *y*_*i*_(*t*) *z*_*i*_(*t*)]^*T*^ be its three-dimensional position within the nucleus, *b*_*i*_(*t*) ∈ {0, 1} its binding state, and *q*_*i*_(*t*) ∈ {0, 1} its activation state (circles in Supplementary Fig. S1A). Particles diffuse freely in the nucleus only when in the unbound state (*b*_*i*_(*t*) = 0); if a molecule is bound (*b*_*i*_(*t*) = 1) it does not move. The diffusion of unbound particles is described by a Brownian motion process by the following stochastic differential equation [57]:

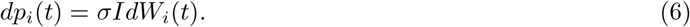

In this equation, *W*_*i*_(*t*) is a standard three-dimensional Wiener process with zero mean and unit rate of variance, and *I* is the identity matrix. The volatility constant *σ* is related to the effective diffusion coefficient *D* through the following equation:

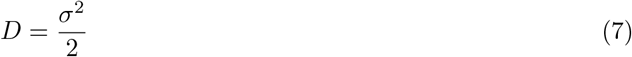

The diffusion coefficient *D* corresponds to the speed of *effective diffusion*, thereby implicitly accounting for otherwise unconsidered biophysical processes, such as protein-protein interactions, collisions with other macromolecules or transient binding to DNA. The diffusion process is assumed to be isotropic and space-homogeneous, as apparent from Eq. (6). Particles are constrained to always stay within the nucleus domain. This boundary condition is not formally accounted for in Eq. (6), but met by a custom reflection scheme described in further detail in section.

In the beginning of the simulation, all particles are unbound and free to diffuse in the nucleus, i.e., *b*_*i*_ = 0 ∀*i* ∈ *P*, where *P* is a set containing all particles. A particle *i* switches from the inactivated (*q*_*i*_(*t*) = 0) to the activated (*q*_*i*_(*t*) = 1) state depending on its presence in the SPB-proximal region of the nucleus and according to an activation probability *P*_*act*_ (asterisk-marked arrow in Supplementary Fig. S1A). The transition of a particle *i* from the unbound to the bound state at time *t* is stochastic and occurs when an activated (*q*_*i*_(*t*) = 1) particle diffuses into the attraction sphere of a *Pre*-state origin (Supplementary Fig. S1C), and according to a binding probability *P*_*bind*_. Particles have to diffuse out of the origin’s attraction sphere before they can try to bind to the same origin again.

In the **origin-centric module** of the model, the progression of the replisome along the DNA sequence following the firing of replication origins is simulated. Each replication origin is associated with a time-dependent, discrete state (circles in Supplementary Fig. S1B) based on the stochastic hybrid model introduced by Lygeros *et al*. [47]. Transitions of replication origin *j* from any state *A* to any state *B* at time *t* are governed by guards 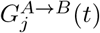.

In the beginning of the simulation, all replication origins are in the pre-replicative (*Pre*) state, meaning that they are licensed but not yet fired. Transitions to the bidirectional replicative (*RB*) state, in which replication is performed in both directions along the DNA sequence, occur when a particle binds to an origin. The origin-centric module is thus closely linked to the particle-centric module, since transitions of origins from the *Pre* to the *RB* state depend on the stochastic hybrid dynamics of the particles:

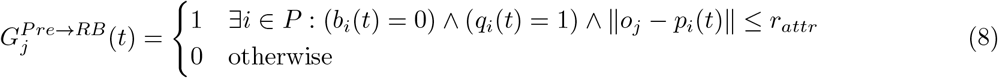

The attraction sphere of an origin is given by the origin’s three-dimensional position *o*_*j*_ and the attraction sphere radius *r*_*attr*_, which is assumed constant and identical for all origins. Only origins in *Pre* state can be bound (fired), and origins can only be bound (fired) once during a replication cycle to prevent re-replication. As mentioned before, to account for the probabilistic nature of the particle-origin binding, the guard 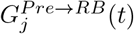 is only evaluated with probability *P*_*bind*_, which is assumed constant and identical for all particles and origins. For simplicity, the probabilistic character of this transition is not formally accounted for in Eq. (8).

Upon origin firing, the bound particle splits into two monomers, with each one associated with one of two newly-formed replisomes. The two replication forks begin to travel along the DNA sequence in opposite directions, thereby marking visited loci as replicated. The fork speed *v*_*fork*_ is assumed constant and identical for all replication forks across the genome [29, 84]. At any time *t*, the base-pair positions 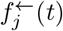 and 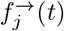 of the two moving replication forks belonging to the fired origin *j* are given by the following equations, where *c*_*j*_ and *t*_*j*_ are the origin’s base-pair position and firing time, respectively:

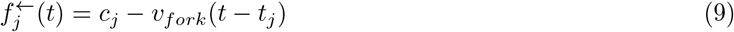

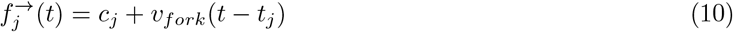

Visited replication origins that are licensed, but have not yet fired, are blocked from further firing attempts (*passive replication*) to prevent re-replication. For a given origin *j*, the transition from the pre-replicative (*Pre*) to the passively replicated (*Pass*) state is governed by the following guard, where the selector *O*_*S*_(*t*) denotes all origins with a state present in set *S* at time *t*:

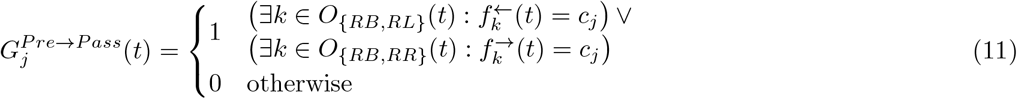

Replication forks travel until they collide with neighboring replication forks or reach the end of their region for which the replication process is simulated (*contig*). For a given fired origin *j*, the first stopping event of a replication fork results in a transition from the bidirectional (*RB*) to a unidirectional (*RL, RR*) replication state, whereas the second collision causes a transition to the post-replicative (*Post*) state:

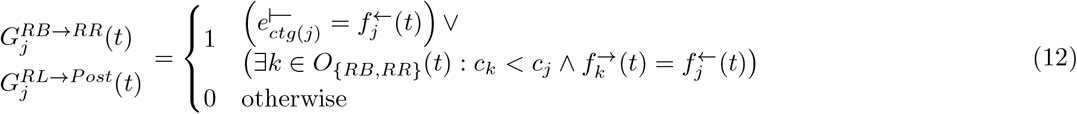

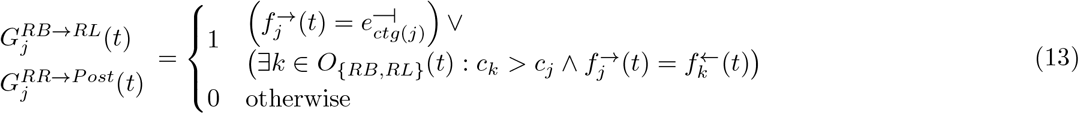

Here, *ctg*(*j*) indicates the contig of origin *j*. The base-pair positions 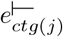 and 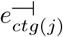 denote the left (5’-end) and right (3’-end) boundaries of the contig, respectively. Upon fork collision, the two colliding monomers join together and form a new firing factor that is released at the collision site. Similarly, to keep the number of firing factors within the nucleus constant during the replication process, for each contig, the pair of replication forks reaching the ends of the respective contig releases a single firing factor as well. As shown in Supplementary Fig. S1D, the particle dimer is released to freely diffuse in the nucleus, triggering its transition back into the unbound state (*b* = 0). In this way, the deterministic dynamics of the origin-centric module affect the stochastic dynamics of the particle-centric module.

### Selection of an ensemble of structures

To simulate our DNA replication model, we used published three-dimensional genome structures derived from coarse-grained polymer modeling [39, 53]. In brief, each coarse-grained polymer model was generated by optimizing a scoring function, which took into account chromosome flexibility, a Rabl-like conformation and chromosomal contacts captured by proximity-ligation from cells synchronized in G1 phase. Using this Monte Carlo approach, a total of 1000 putative genome structures were obtained. The genome structures were further evaluated on their ability to reproduce six sets of fluorescent *in situ* hybridization (FISH) experiments [39].

As input to our model, we selected a representative subset out of the 1000 available structures. This step does not only reduce the computational burden, but additionally increases the trustworthiness of the input data, which are vital for the conclusions made. To select and evaluate the structures, we exploited two publicly available datasets that were not used during the structure generation:

1. **18-pair dataset**: A total of 18 3D Euclidean distances between different gene loci, measured microscopically in a population of unsynchronized fission yeast cells [85].
2. **Cell-cycle dataset**: A set of independent FISH experiments, where two pairs of loci (pair L1 and pair L2) were tracked at various timepoints during the cell cycle [65]. For the purposes of simulating DNA replication, only the following timepoints were considered: early G1, mid G1, end G1, and mid S phase.

We initially assessed the quality of the whole 1000-structure population using both datasets. To examine the agreement with the 18-pair dataset, we measured the minimum pairwise 3D Euclidean distances between all 18 pairs of loci (chromosomal regions) in the 1000 inferred structures and computed the Spearman correlation coefficient (*ρ*) with the FISH distances. As shown in Supplementary Fig. S2A, the average values of the minimum distances across all 1000 structures correlate positively with the FISH data.

However, we observed a large variability when computing the correlation for each structure individually. Specifically, the best-scoring structure exhibited a correlation coefficient of *ρ* = 0.83 (p-value = 6.5 · 10^*−*4^), whereas the correlation was statistically non-significant for the worst-scoring structure (*ρ* = −0.16, p-value = 0.52).

To assess the agreement with the cell-cycle dataset, we computed the minimum distances for the L1 and L2 loci pairs in the 1000-structure population and compared them to the experimental ones. As shown in Supplementary Fig. S2B-C, the corresponding distributions of the inferred and experimental distances exhibited a small overlap, with inferred distances being significantly larger.

Given the above results, we aimed to select an ensemble of structures out of the 1000 available ones that best represent the genomic architecture of a population of fission yeast cells. To achieve this, we employed the following Markov Chain Monte Carlo (MCMC) sampling scheme. We first fitted a multivariate distribution for the 18-pair dataset and two univariate distributions for the L1 and L2 cell cycle dataset using a Bayesian information criterion (BIC). Through sampling from the 1000 structures, we aimed to approximate the distribution of each feature (L1, L2, 18-pairs) separately. For each feature a uniform prior was chosen and the sampler was run for 500 multiple chains with a length of 1000 and a burn-in of 50%. To test the convergence of the multiple chains, the Gelman-Rubin variance test was used. To create the final ensemble, the top 30% of structures with the highest posterior probability distribution for each feature was chosen. This resulted in a final selected ensemble of a total of 217 structures.

Assessing the quality of the selected ensemble of structures indicated a higher agreement with experimental data. As shown in Supplementary Fig. S2D, for the 18-pair dataset, there is an increase in the overall correlation coefficient between the average inferred distances and the FISH distances (*ρ* = 0.73 and *p*-value = 9.5 · 10^*−*4^). At the same time, after the sampling we can approximate the L1 and L2 distances better, since the corresponding distributions shifted to the left and the overlap with the experimental distance distributions increased (Supplementary Fig. S2E-F). Since the sampler was run independently for each feature, we note that not all selected structures agree with each experimental dataset equally well; this explains the remaining discrepancies between the distributions.

### Model discretization and implementation

To evaluate the model within finite time, continuous dynamics of the model need to be discretized. The stochastic diffusion of particles is therefore approximated by a jump-diffusion process as previously described [56], resulting in the gridding of both time and space.

At the beginning of a simulation, particles are initialized randomly within the continuous nucleus domain using rejection sampling, thus defining individual diffusion grids for each particle. The time domain is discretized by sampling random time intervals *τ*_*i*_ from an exponential distribution parameterized with rate parameter *λ*:

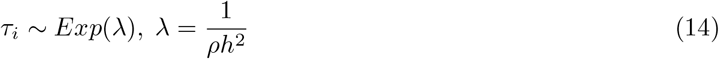

The parameter *ρ* is related to the volatility constant *σ* [86, 87] and thereby to the effective diffusion coefficient *D*:

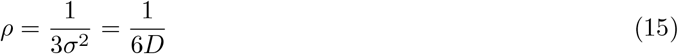

The gridding parameter *h* defines the three-dimensional step size of a particle and is chosen such that particles do not miss replication origin sites by “jumping” over them. It is therefore set to a value smaller than the diameter of the origin attraction sphere:

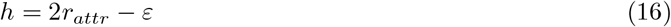

Combining Equations (15) and (16) yields the following rate parameter *λ*:

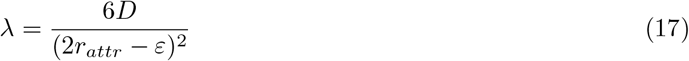

The DNA replication model is implemented in C++11 using the Boost^6^ and Eigen^7^ libraries. MPI^8^ is used to distribute simulations across multiple compute nodes, enabling efficient parameter space exploration.

For the naive implementation, the expected execution time 𝔼[*T*_*exec*_] for the simulation of one full DNA replication cycle is proportional to *T*_*repl*_ · 𝔼[*τ*_*i*_]^*−*1^, where *T*_*repl*_ is the (simulated) time until completion of DNA replication. Plugging in Eq. (17), the proposed algorithm is thus within O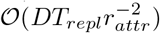:

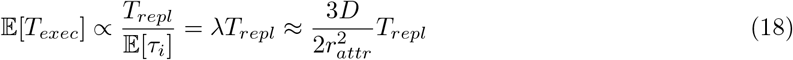

Additional algorithmic optimizations have been implemented to further improve effective simulation speed. In particular, the explicit step-wise simulation of particle diffusion and fork progression dynamics can be skipped for simulated time ranges during which particle diffusion dynamics cannot affect the otherwise deterministic fork progression dynamics (e.g. when all firing factors are bound to a replicating origin).

### Boundary reflection

To ensure that particles always stay within the nucleus domain, they are reflected at the domain boundaries when aspiring to leave the domain. The following reflection scheme for convex boundaries was inspired by Kushner [87]:

1. Let **x**_**asp**_ ∈ ℝ^3^ be the aspired particle position outside the nucleus domain.
2. Define a plane at the position **x**_**asp**_ with the normal **n** = **x**_**asp**_ and move the plane along its normal until it has yielded three candidate points **C** = [**c**_**1**_, **c**_**2**_, **c**_**3**_] ∈ ℝ^3*×*3^ on the particle’s discrete diffusion grid.
3. Solve the linear system of equations **C** · **p** = **x**_**asp**_ to obtain the unnormalized candidate probabilities **p** = [*p*_1_, *p*_2_, *p*_3_]^⊺^. As long as the system does not have a unique real solution (i.e., as long as *rank*(**C**) ≠3 holds true), keep replacing candidate points in **C** by further moving the plane.
4. Randomly draw the reflected particle position **x**_**new**_ from the candidate points **C** according to the unnormalized candidate probabilities **p**.

For boundaries that are not convex, such as the nucleolus boundary, particles are reflected to grid points that are closest to their aspired position (Euclidean distance).

## Quantification and Statistical Analysis

### Radial kernel density

For data analysis, the density of origins of interest at position **x** is estimated as follows:

1. All replication origin positions **x**_*i*_ are rotationally projected to two dimensions:

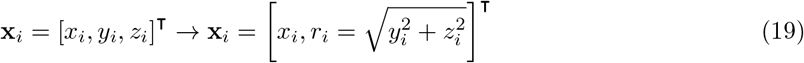
2. For a given set of points {**x** : *i*∈ N, *i*≤*n*, the kernel density estimate 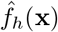 (**x**) with bandwidth *h* at position **x** = [*x, r*]^⊺^ is computed using a two-dimensional Gaussian kernel *K*_*σ*_ with standard deviation *σ* = *h*:

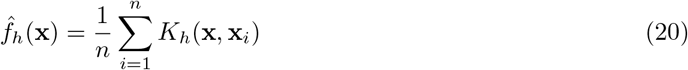

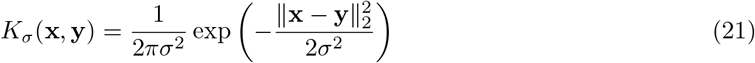
3. The radial kernel density estimate 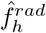 is then obtained by dividing the kernel density estimate 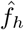 by the corresponding rotational unit volume *V*_1_:

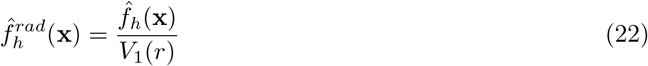

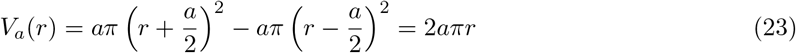

To exclude effects of the underlying replication origin distribution, the radial kernel density estimate of origins of interest was computed relative to the total radial kernel density estimate 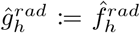 of all origins, where *ε >* 0 is the machine epsilon:

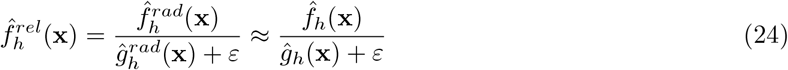

### Categorization of origins for kernel density plots

Regions of efficiency/inefficiency and early/late firing were depicted by plotting the relative radial kernel density estimate of origins whose efficiency or firing time met a certain threshold. Efficiency and firing times were calculated in two different ways. The first was a “structure-averaged” calculation (Fig. 3, Supplementary Fig. S4). Each origin efficiency or firing time was calculated by averaging all simulations of the entire ensemble of structures. The origin was therefore always labeled by the same value when categorizing it as efficient/inefficient or early/late firing. This approach facilitates direct comparison to experimental data, where only the average efficiency or firing time is known for each origin.

In order to visualize the nuances of replication, an “individual structure” calculation was also computed (Supplementary Fig. S3). Based on the set of 5 simulations performed for each structure in the ensemble, the efficiency and average firing time was calculated for each origin. As a result, different values were used to categorize the origin as efficient/inefficient or early/late firing depending on the structure, and origins could be considered, for example, efficient in one structure and inefficient in another. Since the efficiency and firing time of the origins were calculated separately for each structure, this KDE is no longer directly comparable to experiment, but it better captures the details of individual replication profiles.

### Origin firing concurrence

Given a set of completed simulations, origin firing concurrence *C*(*a, b*) indicates competitive relationships between two origins *a, b*. A low concurrence value indicates that the two origins tend to fire together or not at all, whereas a high value indicates that only one of the pair tends to fire at a time. Average efficiencies AvgEff(*a*) and AvgEff(*b*) are used to normalize the count:

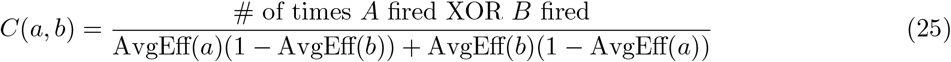

### Replication rate

Replication rate provides insight into the kinetics of DNA replication and the effect of variables such as activation and binding probabilities. The replication rate *R*(*t*) at a given time *t* is calculated by determining the number of active forks *n*_*forks*_(*t*) (two for every origin in *RB* state and one for those in *RL* or *RR* state) and multiplying by the fork velocity *v*_*fork*_:

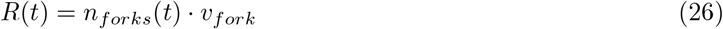

### Probability of firing an origin per unit time and per length of unreplicated DNA

The function *I*(*t*) is a useful characteristic of DNA replication models. Arbona et al. [50] defined the “simulated firing rate per length of unreplicated DNA” as:

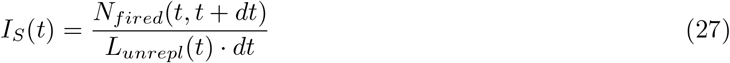

*N*_*fired*_(*t, t* + *dt*) is the number of origins fired between time *t* and *t* + *dt*, and *L*_*unrepl*_(*t*) is the length of unreplicated DNA at time *t*. This function is proportional to *I*(*t*) because the firing rate is proportional to the probability of firing an origin per unit time. To generate *I*_*S*_(*t*) profiles, a boxcar function with width 2 .*dt* and height 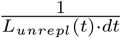 was placed centered at each origin firing time *t. dt* was chosen to be one minute. The sum of all such functions in one simulation run forms *I*_*S*_(*t*) as defined above, and 500 *I*_*S*_(*t*) profiles were averaged for each parameter set to identify general characteristics.

## Experimental Methods

### Growth conditions

Standard fission yeast media and methods were used [88]. All fission yeast strains in this study were grown in supplemented Edinburgh Minimal Medium (EMM) under agitation until mid exponential phase.

### IdU immunofluorescence

In order to achieve IdU incorporation in fission yeast cells, a cell strain (ZP323) expressing the herpes simplex virus thymidine kinase (hsv-TK) and the human equilibrative nucleoside transporter 1 (hENT1) from the constitutive promoter adh1 was employed [89]. To monitor active DNA replication in respect to unclustered centromeric configuration, ZP323 was crossed in a Mis6-GFP, Csi1*Δ* strain (ZP348), originally generated in [64], resulting in ZP370. The isogenic wild type strain (ZP366) was produced by crossing out the Csi1 deletion. In each case, following the crosses hsv-TK and hENT1 inheritance was tested by colony PCR, while Mis6-GFP inheritance was monitored microscopically.

### LacO-LacI strains for 3D distance measurements

For the construction of the cell strains used for SPB-origin 3D distance measurements, the following steps were performed: A modified version (ZP382) of an original strain stably expressing lacI-GFP (ZP255), that was kindly provided by Yoshinori Watanabe, was generated. A PCR-based module method [90] was used to tag the centromeric Mis6 protein at its endogenous locus with the fluorescent protein m-Cherry in ZP382. The pKS391 m-Cherry tagging template was obtained from Kenneth Sawin [91]. The resulting strain (ZP383) was used to generate all lacO-tagged strains. LacO repeats were integrated next to six different origins of interest using a two-step protocol, initially described by Meister et al.[92] in budding yeast, with modified selectable markers for the fission yeast system. Briefly, a hygromycin cassette was amplified from p930 (Addgene #35122) using long primers (100 bases) and guided through sequence homology to the desired integration sites (ZP389-392, ZP396 and ZP401). p991, containing the lacO cassette (kindly provided by Susan Gasser), a ura4+ marker and hphmx4 homologous sequences, was linearized and guided to the already integrated hphmx4 cassette. Valid integrands (ZP418, ZP422, ZP425, ZP429, ZP433, ZP437) were spotted using a double hygromycin and uracil selection process. The boundaries of the integrated lacO cassette were verified by colony PCR.

Origin number, efficiency, genomic position (center of the origin sequence) were based on Heichinger et al. [29], and the respective lacO integration sites are presented for each of the six origins of interest in the following Table:

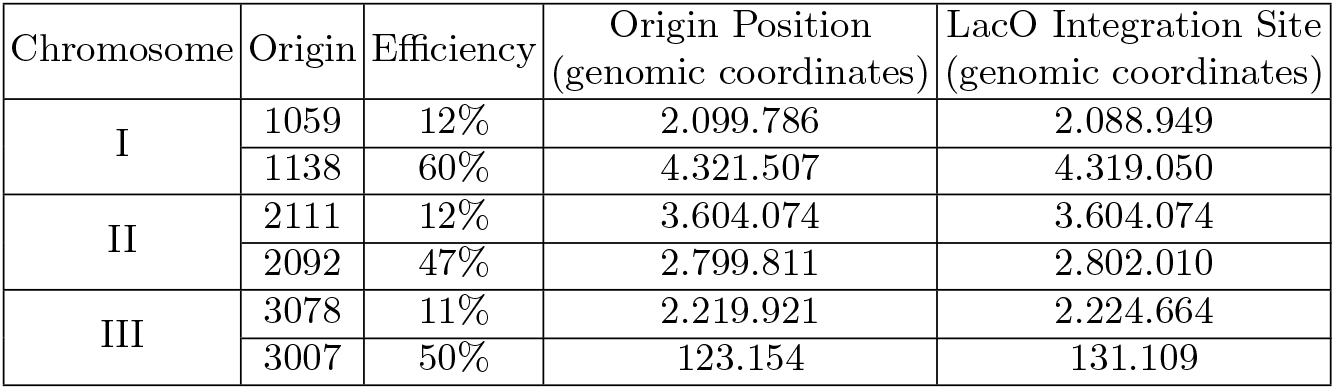

### Real time PCR for efficiency determination

For the determination of origin firing efficiencies, genomic DNA was isolated from an asynchronous population of cells, which consists mostly of cells in the G2 phase of the cell cycle with 2C DNA content [93], and from HU treated cells which are arrested in early S phase. At this stage, origins of replication that have already fired are in two copies per cell, while the rest of the genome is in one copy per cell. Real-time PCR was performed targeting the origin of interest and a dormant origin (ars727) as a control. The efficiency of firing was calculated by dividing the origin/non-origin ratio in HU arrested cells by the origin/non-origin ratio in G2. The exact sequences of the oligonucleotides used for efficiency determination are provided in the Key Resources Table.

### IdU labelling and immunofluorescence

Cells were cultured at 32^*°*^C, in EMM containing the appropriate supplements, to mid-exponential phase (OD∽ 0.2). 500 nM of the thymidine analog IdU and 22 mM hydroxyurea (HU) were simultaneously added in culture. Cells were incubated in the presence of IdU and HU for the duration of a cell cycle (3 hours), fixed in culture by the addition of paraformaldehyde (PFA) in a final concentration of 4% and immunostained against IdU following an immunofluorescence protocol which preserves nuclear morphology, developed by Atanas Kaykov in [62]. Briefly, cells were washed three times in PEM (100 mM Pipes, 1 mM EGTA, 1 mM MgSO_4_, pH = 6.9) and once in PEMS (PEM plus 1.2 M Sorbitol). Cell wall digestion was performed in PEMS containing 0.5 mg/mL Zymolyase-100T and 0.2 mg/ml lysing enzymes for 1 hour at 37^*°*^C. The sample was washed once with PEMS and cells were gradually permeabilized in PEMS containing 0.2% Triton X-100, followed by PEMS containing 0.5% Triton X-100 on ice for 10 minutes. Cells were then washed once in PEM and incubated in PEMBAL (PEM + 1% BSA, 100 mM Lysine hydrochloride, 0.1% NaN_3_, pH = 6.9) containing 5 mM MgSO_4_ for 30 minutes under rotation. Incorporated IdU was exposed by incubating cells for 2 hours at 37^*°*^C in 100*μ*l PEMBAL/5 mM MgSO_4_, containing a 1:2 dilution of nucleases from the 5-Bromo-2′-deoxy-uridine labeling and detection kit I (Roche), and a 1:10 dilution of mouse anti-BrdU antibody (BD Biosciences, 347580). Cells were washed three times in PEMBAL/5 mM EDTA, recovered in 100 *μ*L PEMBAL containing 1:10 mouse anti-BrdU (from BD) and incubated on a rotating wheel overnight. Cells were then washed three times in PEMBAL, incubated with Alexa Fluor 568 goat anti-mouse antibody (Invitrogen, A-11031) for 3 hours, washed in PEMBAL three times, then once in PBS, stained with DAPI and mounted. Imaging was performed using a widefield Olympus microscope.

### Live imaging for 3D distance measurements

Cells were grown in supplemented EMM to at 25^*°*^C until an OD of approximately 0.2 (4 10^6^ cells/mL, mid-exponential phase). Cell growth in lower temperature improved the signal to noise ratio for the lacO-lacI dots. Glass-bottomed imaging plates (Ibidi) were coated with lectin from Glucin max (2 mg/mL) for 10-15 minutes. Lectin was removed and plates were left to dry. 1 mL of the cell culture was added to the lectin coated plate, and cells were allowed to sediment on the plate for 10 minutes. Excess medium was removed, and 1 mL of pre-heated filtered medium was gently added on top of the cells.

Stable temperature conditions for live imaging were achieved using a temperature-controlled microscopy chamber. The acquisition parameters were optimized to minimize phototoxicity. To determine 3D distances, imaging was performed at multiple optical sections using a Leica SP5 confocal microscope. 3D distances were measured using the ImageJ “Boris” plug-in.

### Data and Code Availability

The source code of the DNA replication model and all generated *in silico* data are available under an open source license at: https://github.com/AI4SCR/dna-replication.

## Supplementary Tables

**Table S1:** Model parameters and inputs for all simulation runs.

|  | **Model variant 1:**<br>**Random origin**<br>**positioning** | **Model variant 2:**<br>**Realistic origin**<br>**positioning** | **Model variant 3:**<br>**SPB-mediated particle**<br>**activation** |
| --- | --- | --- | --- |
| Origin Positioning | 100 random structures [53,<br>“confined” model],<br>randomly selected | 217 experimentally derived structures [53,<br>“interactions” model], selected using MCMC<br>(Supplementary Methods ) |  |
| Particle activation | Already active |  | Activated in the SPB |
| Origin Coordinates | 893 origins (base pair coordinates) [29] |  |  |
| Fork speed | 50 bp/s [29] |  |  |

| **First simulation run** |  |  |
| --- | --- | --- |
| Number of particles ($N$) | {50, 100, 150, 200, 400, 600, 800, 1000} | |
| Attraction radius ($\mu m$) | {0.015, 0.025, 0.035} | |
| Effective diffusion coefficient ($\mu m^2/s$) | {0.5, 2, 3.5} | |
| $P_{bind}$ | 1 | |
| $P_{act}$ | Not relevant | 1 |

| **Second simulation run** |  |  |
| --- | --- | --- |
| Number of particles ($N$) | Not simulated | {150, 200, 250, 300, 350,<br>400, 450, 500} |
| Attraction radius ($\mu m$) | | 0.025 |
| Effective diffusion coefficient ($\mu m^2/s$) | | 0.5 |
| $P_{bind}$ | | {0.1, 0.01, 0.001, 0.0005} |
| $P_{act}$ | | {0.01, 0.001, 0.0003,<br>0.0001} |

## Supplementary Figures

**Fig. S1:**
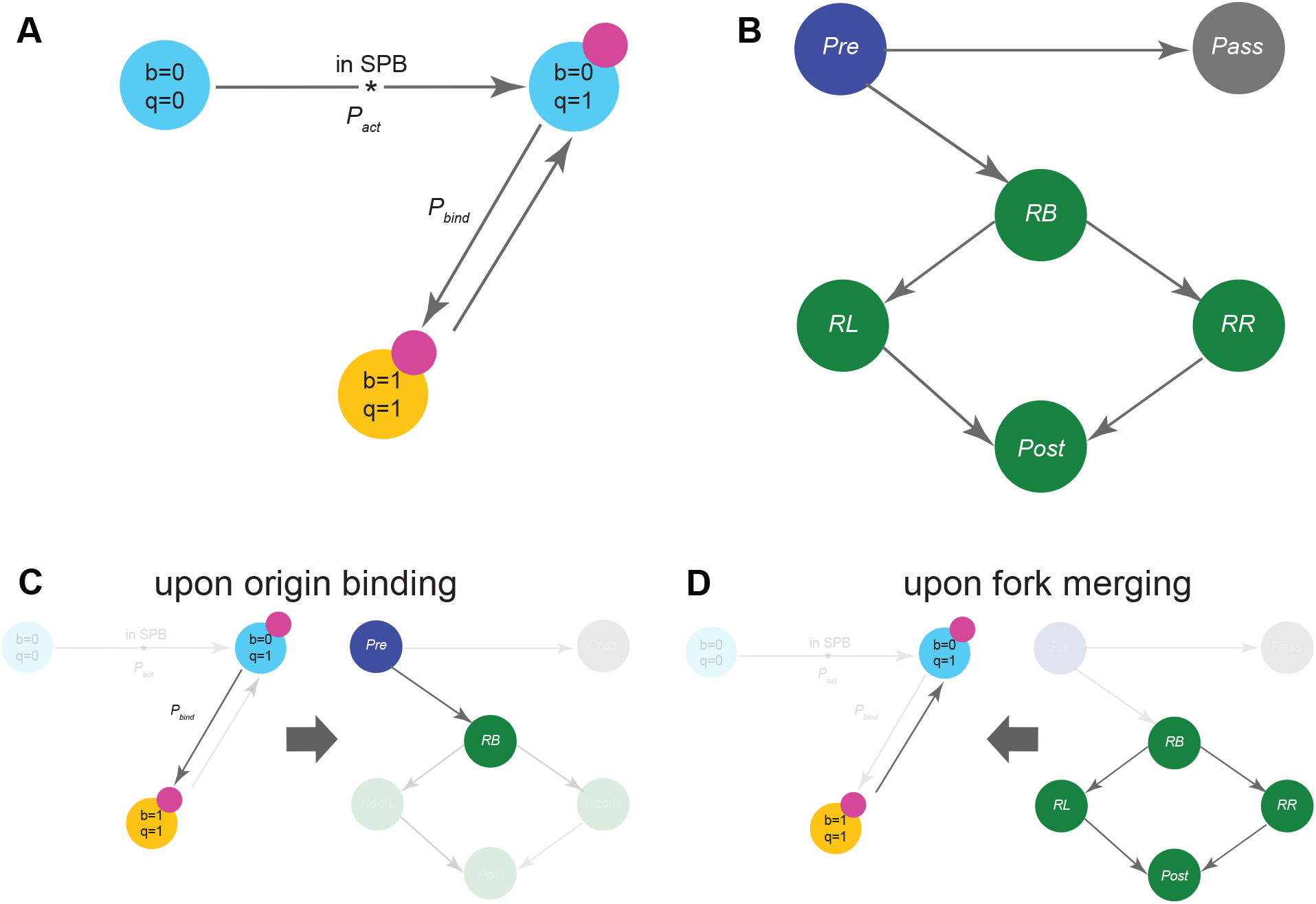
(A-B): Overview of model states and transitions. All discrete states (circles) and allowed transitions (arrows) of the particle-centric (A) and origin-centric (B) modules of the model. Particle and origin states are shown using the same colors as in Fig. 1. Transitions to the activated and bound particle states are probabilistic (model parameters *P*_*act*_, *P*_*bind*_). The asterisk indicates a transition present only in the SPB-mediated activation variation of the model. **(C-D): Crosstalk between the two modules of the model. (C)** When an activated (*q* = 1) and unbound (*b* = 0) particle diffuses into the attraction sphere of an origin, it binds to it with probability *P*_*bind*_ and undergoes the *b* : 0 → 1 transition. This causes the origin to undergo the *Pre* → *RB* transition and initiate firing. These transitions are stochastic and depend on the continuous dynamics of the model (particle diffusion). **(D)** When replication forks collide, the corresponding origins undergo the *RB* → *RL* | *RR* or *RL* | *RR* → *Post* transitions. The latter causes the colliding firing factor monomers to dimerize, releasing a new particle into the nucleus (*b* : 1 → 0 transition), which is free to diffuse and potentially fire other origins.

**Fig. S2:**
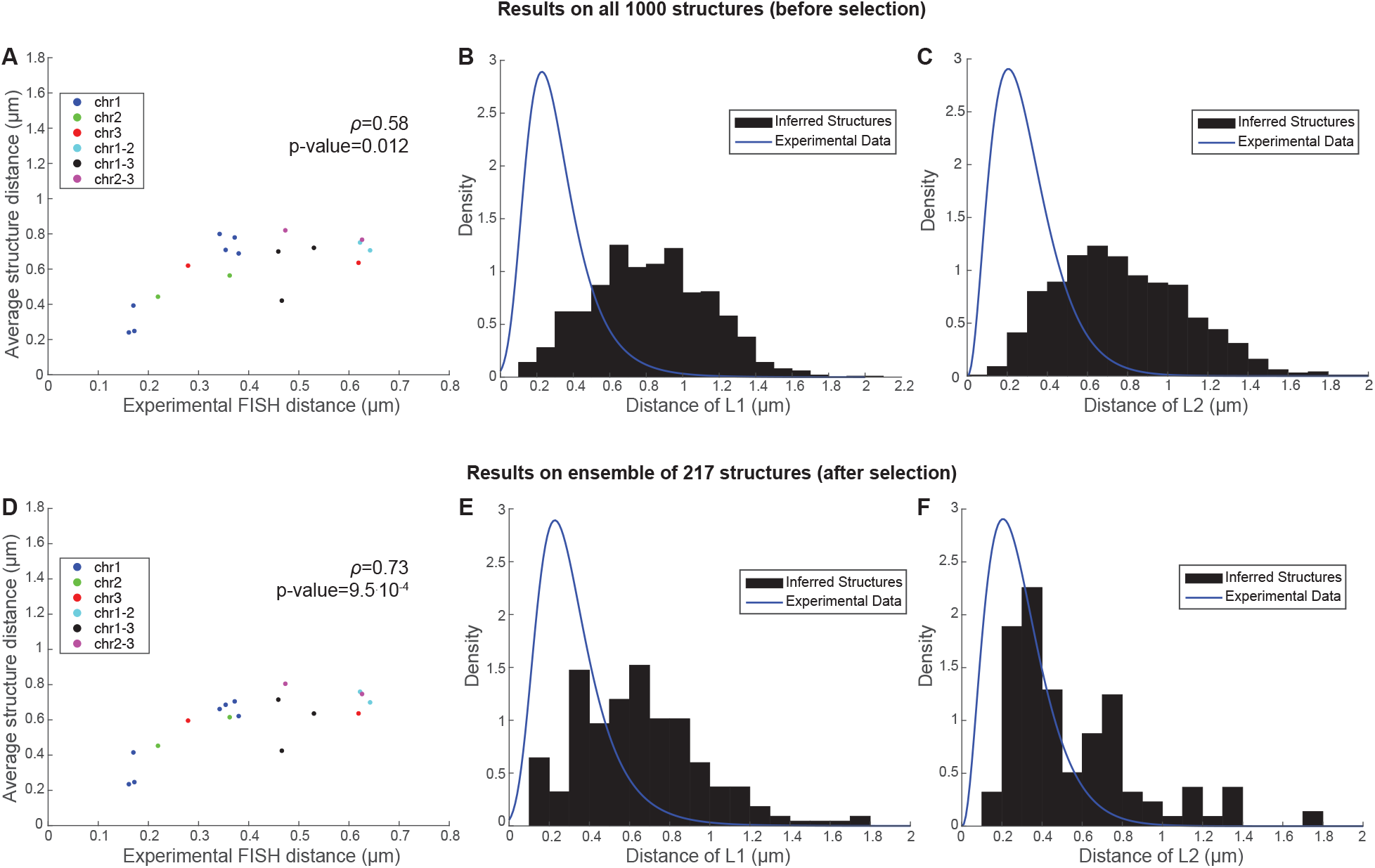
Selection of a representative ensemble of structures. (A-C) Comparison of the total 1000 structures with experimental datasets. (A) Correlation between the 18-pair FISH measurements and average 3D Euclidean distances, as computed from all 1000 inferred structures. Each dot represents a distance in an intra- or inter-chromosomal loci pair (colors given in legend). (B-C) Distribution of population of structures for the L1 (B) and L2 (C) loci pairs. Blue represents the model of the probability density, fitted to the corresponding experimental data that minimizes the Bayesian Information Criterion. Black histograms represent the distribution of the specific distances in the whole 1000-structure population. (D-F) Same as in (A-C), but for the selected ensemble of structures.

**Fig. S3:**
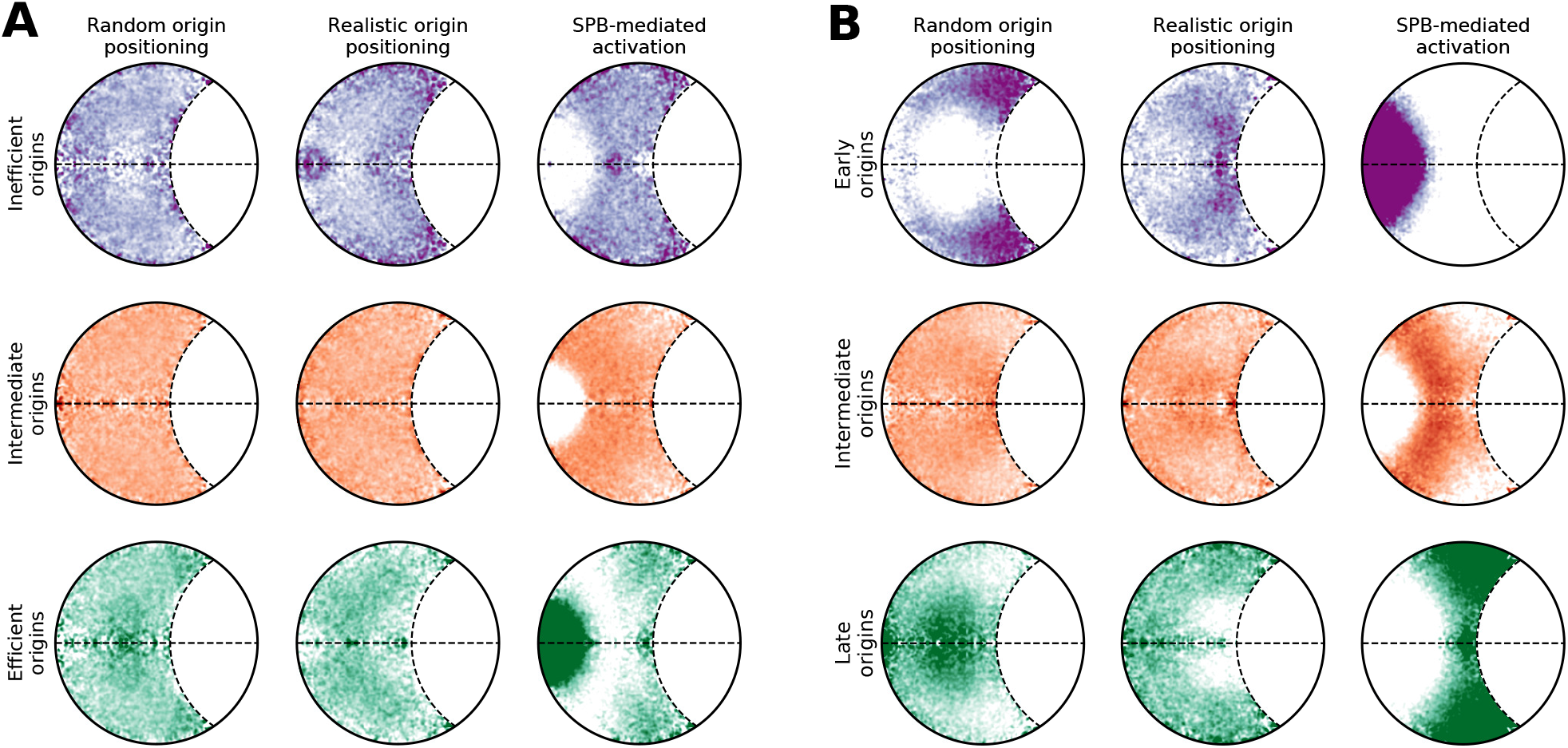
Spatial organization of average origin efficiency and firing times for individual structures. Relative radial kernel density plots were computed as explained in Supplementary Methods Section, “individual structure”. (A) Relative radial kernel density estimates of inefficient (lower quartile efficiencies, top row), intermediate (*Q*_1_ *<* efficiency *< Q*_3_, middle row), and efficient (upper quartile efficiencies, bottom row) origins. (B) Relative radial kernel density estimates of early-firing (lower quartile firing times, top row), intermediate (*Q*_1_ *<* firing time *< Q*_3_, middle row), and late-firing (upper quartile firing times, bottom row) origins.

**Fig. S4:**
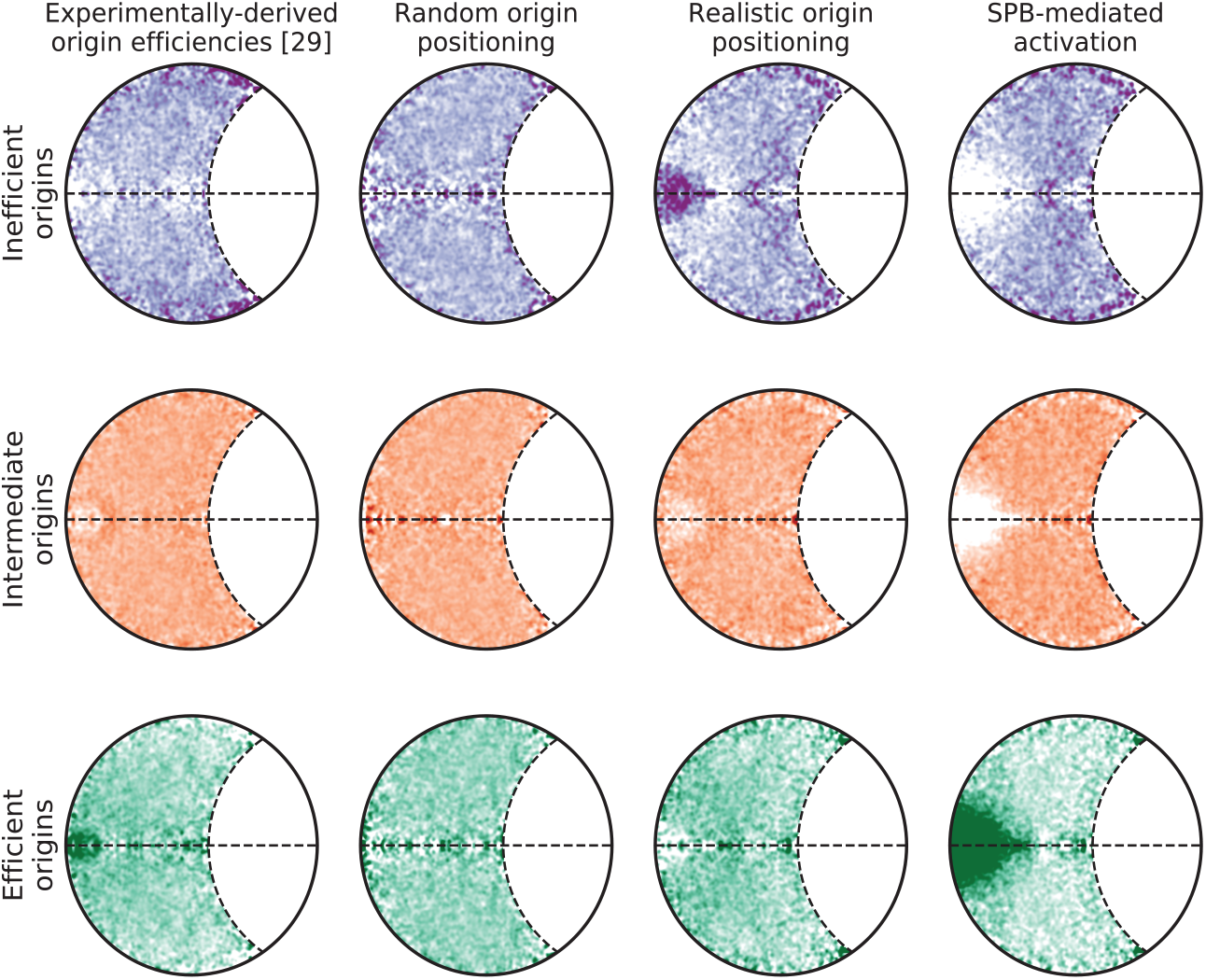
Spatial organization of average origin efficiencies. Relative radial kernel density estimates of inefficient (lower quartile efficiencies, top row), intermediate (*Q*_1_ *<* efficiency *< Q*_3_, middle row), and efficient (upper quartile efficiencies, bottom row) origins for experimental data and all three model variants.

## Footnotes

6 Boost 1.61.0 (http://www.boost.org)

7 Eigen 3.3.4 (http://eigen.tuxfamily.org)

8 MPI 3.1 (http://www.mpi-forum.org)

